# Sustained Maternal Cortisol Elevation Links Metabolic Stress to Fetal Growth Restriction in a Guinea Pig Model of Undernutrition

**DOI:** 10.64898/2026.10.17.759797

**Authors:** Courtney Ruggiero, Priscilla Radillo, Baylea Davenport, Helen Jones, Rebecca Wilson

## Abstract

Fetal growth restriction (FGR) resulting from maternal undernutrition is commonly attributed to insufficient nutrient substrate delivery to the fetus. However, the contribution of maternal physiological stress responses to this outcome remains poorly characterized. Using the well-established guinea pig maternal nutrient restriction (MNR) model, we conducted a comprehensive retrospective analysis to interrogate the hypothesis that maternal stress physiology actively mediates fetal growth outcomes beyond simple substrate limitation. Maternal salivary and serum cortisol were significantly elevated in MNR dams throughout pregnancy despite no baseline differences at conception, indicating that hypothalamic-pituitary-adrenal (HPA) axis activation was specifically triggered by the combined physiological demands of pregnancy and nutritional constraint. The sustained cortisol elevation was accompanied by acute metabolic perturbations at mid-pregnancy, including hypoglycemia and depleted circulating lipids. While most acute metabolic disturbances resolved by near-term, cortisol remained elevated, and MNR fetuses exhibited parallel cortisol elevation alongside sustained growth restriction. Maternal weight divergence emerged only in late gestation and was driven primarily by reduced conceptus weight. Maternal organ analysis revealed selective hepatic enlargement with apparent lipid accumulation in MNR dams near-term. Overall, these findings help to reframe nutrition-related FGR as a condition in which maternal adaptive stress responses, reduced nutritional components and placental insufficiency are mechanistically intertwined, with important implications for the design of effective interventions beyond simply nutrient supplementation.

**New and Noteworthy:** This study demonstrates that maternal physiological stress, not substrate limitation alone, is a driver of impaired fetal growth in a guinea pig model of undernutrition in pregnancy. We show that maternal hypothalamic-pituitary-adrenal axis activation produces sustained cortisol elevation throughout pregnancy independent of nutritional recovery. This stress-related response was accompanied by selective hepatic and metabolic adaptations.

## INTRODUCTION

Fetal growth restriction (FGR) is associated with substantial morbidity and mortality to mother and baby both immediately and across the lifespan. Beyond contributing to over 2.5 million stillbirths annually and increasing preterm birth risk^1–5^, those born growth restricted face elevated rates of cognitive deficits, cardiovascular disease, metabolic syndrome and obesity^6, 7^. Maternal undernutrition, whether from food insecurity, poor quality foods or hyperemesis gravidarum, represents a major extrinsic cause of FGR globally^8, 9^. While the association between maternal nutritional status and fetal growth has been well established through epidemiological studies^10, 11^, the mechanistic understanding linking maternal undernutrition to impaired fetal development remains incomplete. Currently, there is no effective treatment for FGR beyond early delivery and intensive neonatal care^12, 13^, highlighting the critical need to better understand the pathophysiology underlying nutrition-related growth restriction.

The prevailing mechanistic framework for understanding how maternal undernutrition leads to FGR has focused primarily on limited nutrient substrate availability and placental insufficiency^14, 15^. Reduced maternal nutrient intake directly limits the supply of glucose, amino acids and lipids available for placental transfer to the fetus^16^. Additionally, chronic nutrient restriction can impair placental development and function, reducing placental size, altering villous architecture, decreasing vascularity and compromising nutrient transporter expression^17^. This placental insufficiency, defined by the placenta’s failure to transfer adequate nutrients and oxygen for proper fetal growth^18–20^, creates a direct constraint on fetal growth potential. Animal models of maternal undernutrition have consistently demonstrated reduced placental efficiency, decreased expression of nutrient transporters such as GLUT and System A amino acid transporters and altered production of placental hormones^21–25^. While this substrate-focused framework has advanced our understanding of nutrient-restricted FGR, it largely frames the mechanisms of poor fetal growth as due to limited maternal resources. Given that nutritional supplementation in pregnancy often fails to substantially improve fetal growth outcomes^26^, this suggests that nutrient availability alone may not fully explain growth restriction in maternal undernutrition.

Pregnancy itself represents a profound physiological challenge, requiring substantial maternal metabolic adaptation to support both maternal homeostasis and fetal development^27^. Normal pregnancy is characterized by progressive insulin resistance, increased cardiac output, expanded blood volume, altered renal function and significant changes in lipid and glucose metabolism^28^. These adaptations become even more pronounced when maternal nutrient availability is limited, triggering maternal stress responses that extend beyond simple substrate limitation^29^. The maternal hypothalamic-pituitary-adrenal (HPA) axis is a key regulator of stress responses, and activation results in elevated glucocorticoid production^30^. Glucocorticoids play critical roles in maternal metabolic adaptation, mobilizing energy stores through hepatic gluconeogenesis and lipolysis, but chronic elevation can have detrimental effects on both maternal organ function and placental development ^31, 32^. Maternal stress and elevated cortisol have been associated with adverse pregnancy outcomes including FGR^33^. However, the extent to which maternal physiological stress responses contribute to fetal growth outcomes in undernutrition beyond direct substrate limitation remains poorly understood.

We hypothesized that the reduced fetal growth characteristic of maternal undernutrition models reflects not only limited nutrient substrate availability, but also the maternal physiological stress response to meeting the increased metabolic demands of pregnancy. Specifically, we propose that maternal adaptations, including HPA axis activation and altered partitioning of limited nutrient resources, actively contribute to the development of placental insufficiency and FGR. This perspective shifts the mechanistic understanding from a purely substrate-limited model to one in which maternal stress responses contribute to the mediation of fetal outcomes.

To test this hypothesis, we conducted a comprehensive characterization of maternal physiological adaptations to nutrient restriction across pregnancy in the guinea pig. The guinea pig offers advantages for studying maternal-fetal interactions due to similarities with human pregnancy^34^. Unlike mice and rats, guinea pigs exhibit comparable hormonal profiles to humans during pregnancy, produce cortisol rather than corticosterone as their primary glucocorticoid, and have a relatively long gestation period (∼65-70 days) that allows detailed study of temporal changes in maternal physiology ^35, 36^ ^37^ ^30^. Using the well-established maternal nutrient restriction (MNR) model in the guinea pig ^21, 38–40^, we examined maternal responses at two gestational windows: mid-pregnancy (GD 33-38) and near-term (GD 57-63). We measured longitudinal maternal weight gain trajectories, salivary and serum cortisol across pregnancy, maternal blood nutrients, metabolites and hormones, and fetal growth outcomes. This retrospective analysis of control and nutrient-restricted dams allowed us to characterize the progressive maternal physiological response to chronic undernutrition and identify potential stress-related mechanisms contributing to fetal growth restriction beyond direct nutrient limitation.

## MATERIALS AND METHODS

This study represents a retrospective analysis of control and maternal nutrient restricted (MNR) dams collected between 2018-2023 as part of studies investigating the use of *IGF1* gene therapy for the treatment of fetal growth restriction^21, 40^. Only animals that did not receive gene therapy treatment (Control and MNR dams with sham-treated litters) were included in this manuscript.

Animal care and usage was approved by the Institutional Animal Care and Use Committees at Cincinnati Children’s Hospital and Medical Center (Protocol number 2017-0065) and University of Florida (Protocol number 202011236). Dams (Dunkin-Hartley) were purchased at 500-550 g (7-8 weeks of age) and allowed to acclimate to the animal facilities for two weeks prior to study initiation. After the acclimatization period, dams were weighed, ranked from heaviest to lightest, and systematically assigned to either ad libitum diet (termed Control) or MNR diet so that the mean weight was no different between the Control and MNR groups. For both Control and MNR dams, water was provided ad libitum. Food (LabDiet diet 5025: 27% protein, 13.5% fat, and 60% carbohydrate as % of energy) intake in the MNR dams was restricted to 70% per kilogram body weight of the Control group from at least 4 weeks preconception through to mid-pregnancy (GD30). From GD30, food was restricted to 90% per kilogram body weight of the Control group to prevent fetal demise and pregnancy loss^39^. Control dams were provided food ad libitum. Dams were weighed weekly throughout the study period and on gestational day 1 (GD1).

Dams sacrificed at mid-pregnancy (GD33-38; Control: n=6, MNR: n=5) were anesthetized once at GD28-33, and a single intra-placental sham nanoparticle injection was administered via ultrasound guidance^21^. Dams sacrificed at near term (GD56-63; Control: n=7, MNR: n=7) were anesthetized and received intra-placental sham nanoparticle injections three times, 8 days apart (GD36±3, GD44±3, and GD52±3)^40^. All dams were sacrificed by carbon dioxide asphyxiation followed by exsanguination and removal of the heart. Maternal blood was collected by cardiac puncture into blood tubes containing no coagulants and serum obtained by centrifugation. Maternal organs (heart, lung, liver, kidney, spleen, ovary) were excised and weighed. Individual fetuses and maternal-fetal interface tissues (placenta and sub-placenta/decidua) were dissected, counted and weighed. Resorption sites were recorded. Maternal liver and adrenal tissues were processed for histology (fixed in 4% PFA). To ensure rigor and reproducibility, all samples were analyzed in a blinded manner.

### Maternal Saliva and Cortisol Analysis

Maternal saliva samples were collected at the initiation of dietary treatment (approximately 4 weeks preconception), gestational day 1 (GD1), and at gestational days 35, 43, 51, and 58. All saliva samples were collected between 9:00 and 10:00 AM to control for diurnal variation in cortisol levels. To collect saliva, cotton buds were inserted into the cheek pouch, parallel with the cheek teeth for at least 2 minutes^41^. Tips were withdrawn, buds were cut off and centrifuged at 12,500 × g for 2 min to extract saliva. Samples were stored at -80°C until analysis. Salivary and serum cortisol concentrations were measured using the DetectX® Cortisol ELISA Kit (Arbor Assays) according to manufacturer’s protocol and run in duplicate. Concentrations were calculated from a standard curve.

### Serum Metabolite and Hormone Analysis

Serum glucose, electrolytes (sodium, potassium, chloride, calcium, carbon dioxide (CO_2_)), kidney function markers (blood urea nitrogen (BUN), creatinine), liver enzymes (alkaline phosphatase (ALP), alanine aminotransferase (ALT), aspartate aminotransferase (AST), total bilirubin), proteins (albumin, total protein) and lipids (cholesterol, triglycerides and VLDL) were measured using the Abbott Piccolo Xpress Chemistry Analyzer with the Comprehensive Metabolic Panel (CMP) or Lipid Panel reagent discs according to manufacturer’s protocol. Serum lactate was measured using the YSI 2900 Biochemistry Analyzer (YSI Life Sciences) according to manufacturer’s protocol. Serum creatine was measured using the EnzyChrom™ Creatine Assay Kit (BioAssay Systems) according to manufacturer’s protocol.

Serum progesterone, estriol, DHEA-s and oxytocin were measured in serum samples using the DetectX® ELISA Kits from Arbor Assays following manufacturers protocol. Serum insulin was measured using Novus Biologicals Guinea Pig Insulin ELISA Kit following manufacturers protocol. Details of sample dilution and the sensitivity and intra-assay precision reported by the manufacturer can be found in Supplemental Table 1. All ELISAs were performed in duplicate, and concentrations calculated from standard curves.

### Histology and Immunohistochemistry

Paraffin-embedded maternal liver and adrenal tissue sections (5 µm thick) were obtained for histological and immunohistochemical analysis. Sections were dewaxed and rehydrated following standard protocols. All stained sections were imaged on the Axioscan Scanning Microscope (Zeiss), visualized using the Zen Imaging Software (Zeiss) and analyzed using ImageJ software (v1.53a).

#### Maternal Liver

Hepatic lipid accumulation was assessed using standard Masson’s trichrome staining. Formal quantification of lipid accumulation was not performed due to limited sample availability; sections were visually evaluated for the presence of cytoplasmic lipid vacuoles.

#### Maternal Adrenal

Masson’s trichrome staining was used to identify and measure cortical and medullary areas in the Zen Imaging software. Collagen deposition was visualized using Sirius Red staining. Images (20X magnification) of 10 separate fields within the zona fasciculata per section were obtained. The Color Deconvolution plugin in ImageJ was used to split images into red, green and blue channels. With the blue channel image selected, percent area was measured using the Threshold and Measure tools. An average percent collagen was then calculated across the 10 images per section. Cell hypertrophy was assessed by obtaining 10 images (20X magnification) per section from hematoxylin-stained sections. Blue nuclei were separated using the Color Deconvolution tool in ImageJ. The Threshold function was then used to select nuclei, the Watershed tool was used to separate nuclei where appropriate and the Measure Particles tool was used to count the number of nuclei within the images. Cells per mm^2^ was calculated and average across the 10 images obtained per section.

### Statistics

All statistical analyses were performed in R (v4.6.0) using the lme4, lmerTest, emmeans, and tidyverse packages. Distribution assumptions were checked with Q-Q plots. Statistical significance was set at p ≤ 0.05. Data are presented as estimate marginal mean ± SEM, unless otherwise specified.

Baseline characteristics (maternal weight at GD1 and litter size) were compared using Welch’s t-tests. Associations between maternal weight at GD1 and litter size were assessed using Pearson’s product-moment correlation. Fetal sex ratios were compared between diet groups using Fisher’s exact test.

Litter size was included as a covariate for maternal weight gain trajectories after mid-gestation given the established biological relationship between fetal load and maternal weight. For maternal metabolites, litter size was included where associations showed at least a trend-level relationship (p < 0.10).

Maternal weight gain and salivary cortisol trajectories across pregnancy were assessed using linear mixed-effects models with diet, time and their interaction as fixed effects. Animal ID was included as a random effect to account for repeated measures. Post-hoc pairwise comparisons between diets at each week were conducted using estimated marginal means with Bonferroni correction. To determine when the litter size effect on weight gain emerged, sequential analyses were conducted at two-day intervals throughout gestation, fitting linear mixed-effects models with gestational day and litter size as predictors and animal ID as a random effect.

Maternal organ weights, blood metabolites and adrenal morphology variables were analyzed using ANCOVA with diet as the main factor and gestational day and litter size as covariates.

Fetal and placental outcomes were analyzed using linear mixed-effects models with diet, fetal sex and their interaction as fixed effects, and animal ID as a random effect to account for clustering within litters.

## RESULTS

Baseline characteristics at GD1 were assessed in both the mid-pregnancy (n=11) and near-term (n=14) collected cohorts. Data from both cohorts were combined for baseline correlation analyses (total n=25), as weight at GD1 and litter size did not differ between cohorts (both p>0.05). Longitudinal weight gain analyses used only the near-term cohort (n=14), as full pregnancy trajectories were required.

In Control animals, maternal weight at GD1 was positively associated with litter size (r=0.61, R²=0.38, p=0.059). In contrast, MNR dams showed no detectable relationship between baseline weight and litter size (r=0.25, R²=0.06, p=0.44), likely due to compressed litter size variation (range: 2–4 pups) relative to controls (range: 2–5 pups). Exploring further and testing whether metabolic constraints due to MNR created a ‘ceiling effect’ on litter size, we repeated the analysis on data available from 70 dams collected across various other studies the lab performed. In n=34 Control and n=36 MNR dams, viable litter size was not different depending on diet (mean Control: 3.06 vs. mean MNR: 2.81, p>0.05) and documented resorptions rates were low in both Control and MNR pregnancies (Control: 5.9% (2/34) vs. MNR: 11.1% (4/36)). Maternal weight at GD1 predicted litter size in both diet groups (Control: r=0.54, p=0.007; MNR: r=0.33, p=0.05). The relationship appeared descriptively weaker in MNR dams, but this difference was not statistically significant (interaction p=0.51), indicating that dietary restriction did not fundamentally alter the relationship between maternal body size at the time of conception and reproductive capacity.

Sequential analysis revealed that the litter size effect on maternal weight gain emerged progressively during the second half of pregnancy, consistent with accelerating fetal growth. The effect became marginally significant around gestational day 52 (∼75% of the ∼70 day gestation period; β = 18.3 g per pup, p = 0.051. Supplemental Figure 1) and reached conventional statistical significance by day 56 (β = 18.2 g per pup, p = 0.047).

Maternal weight gain increased linearly across pregnancy in both diet groups (p<0.001; Figure 1). After controlling for litter size, there was no significant difference between Control and MNR dam weight in early pregnancy (weeks 1–2, both p>0.0.05; Supplemental Table 2). A trend toward divergence emerged at week 3 (18.5 g difference, p=0.071), with Control dams gaining significantly more weight than MNR from week 4 onward (Figure 1). This difference increased progressively throughout pregnancy at an average rate of 3.5 g per week, reaching 39.7 g difference by week 9 (95% CI: [8.0, 71.4], p=0.016).

**Figure 1.**
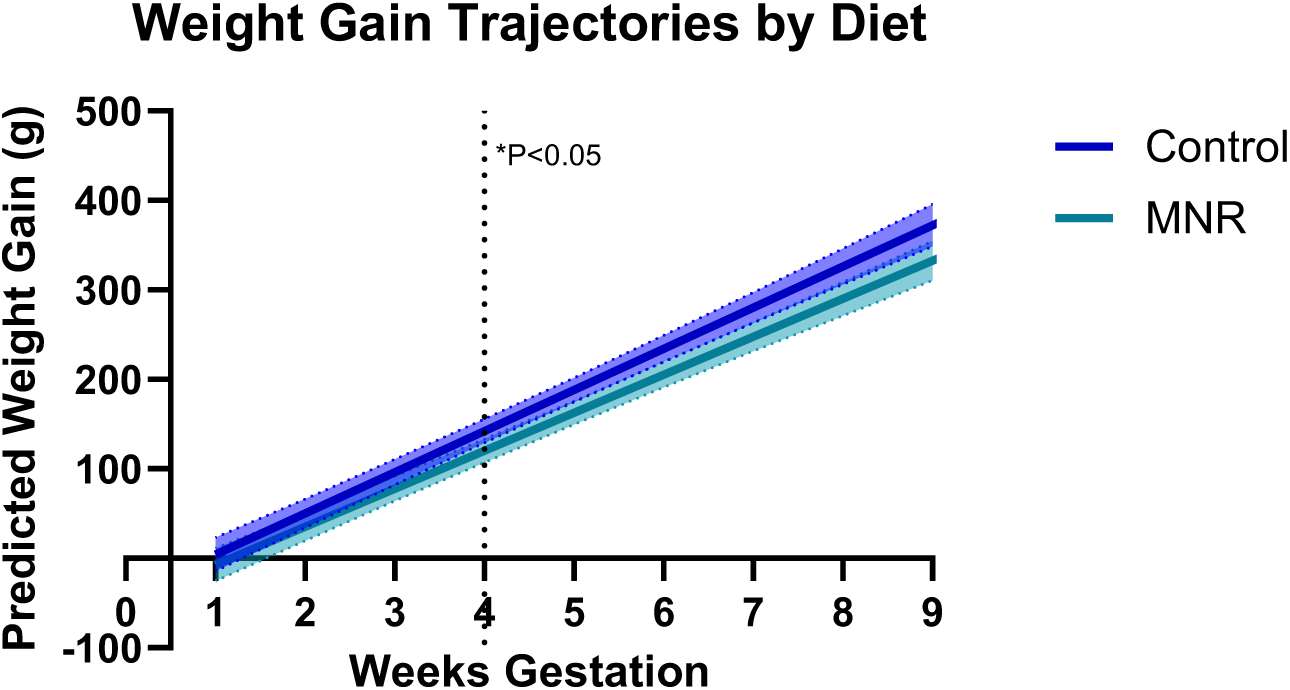
Maternal weight gain in pregnancy comparing maternal nutrient restricted (MNR) dams to ad lib fed Control dams. Maternal weight gain increased linearly across pregnancy in both diet groups, with Control dams gaining significantly more weight than MNR dams from week 4 of gestation onwards. Data are model-predicted weight gain from baseline (g) across gestation for Control (blue) and MNR (green) diet groups, controlling for litter size. Shaded regions represent 95% confidence intervals. The dotted vertical line indicates the first gestational week at which diet reached statistical significance (p < 0.05). Data from term cohort only (n=14: 7 Control, 7 MNR)

### Saliva Cortisol

There was no significant relationship between days on the MNR diet before pregnancy and pre-pregnancy cortisol concentrations (r = -0.13, p = 0.527, n = 26 timepoints from 7 dams; Figure 2A). Linear regression confirmed that diet duration did not predict cortisol levels (β = -0.037, SE = 0.058, p = 0.527, R² = 0.017).

**Figure 2.**
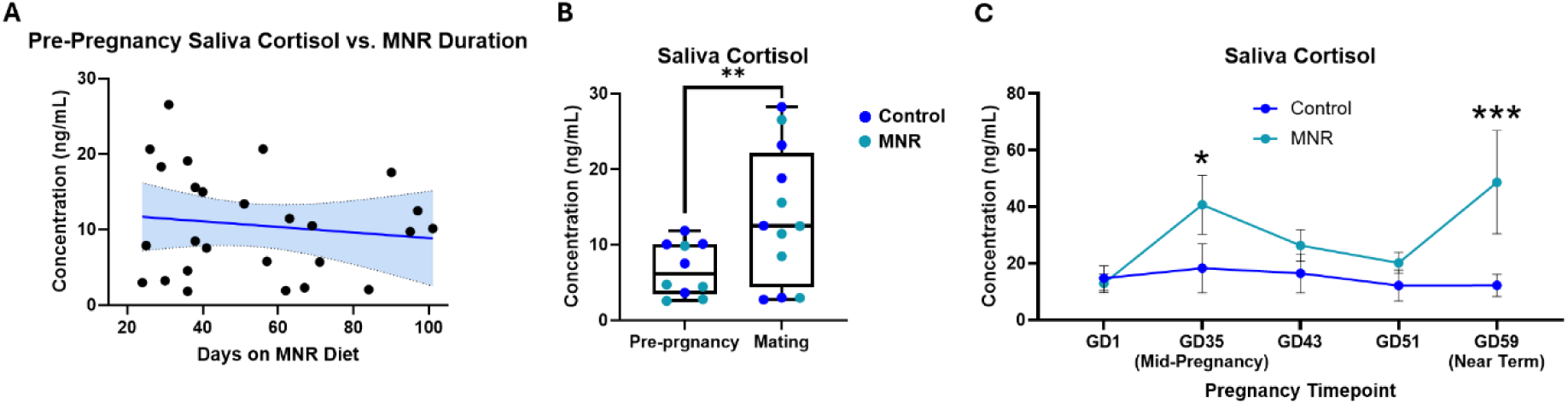
Salivary cortisol concentrations in Control and maternal nutrient restricted (MNR) dams across periconception and pregnancy. (A) There was no significant relationship between days on the MNR diet before pregnancy and pre-pregnancy cortisol concentrations. (B) Mating significantly increased salivary cortisol in both Control and MNR dams when compared to basal salivary cortisol collect at the end of the 4 week dietary conditioning period (pre-pregnancy). (C) MNR animals had elevated basal cortisol compared to Controls at GD35/mid-pregnancy and GD59/near term. Data are: (A) pre-pregnancy salivary cortisol as a function of duration on the MNR diet. The solid line represents a linear regression fit with 95% confidence interval (shaded); (B) median and interquartile range with individual data points overlaid; (C) estimated marginal mean ± SEM, calculated from linear mixed effects modelling. Asterisks denote significant between-group differences (*p < 0.05, **p < 0.01, ***p < 0.001). n=14: 7 control, 7 MNR

Mating significantly increased salivary cortisol in both Control and MNR dams when compared to basal salivary cortisol collect at the end of the 4 week dietary conditioning period (p = 0.001; Figure 2B). Baseline analysis of saliva cortisol at mating/GD1 showed no difference between Control and MNR dams (Control: 14.8 ± 4.3 vs. MNR: 12.9 ± 3.2 ng/ml, p=0.682). Analysis of saliva cortisol trajectories across pregnancy revealed a significant effect of diet (p = 0.018) and marginally significant diet and pregnancy timepoint interaction (p = 0.088). Post-hoc comparisons revealed that MNR animals had elevated basal cortisol compared to Controls at GD35/mid-pregnancy (Figure 2C) and GD59/near term (Figure 2C). Blood cortisol was also elevated in MNR dams compared to Control at these pregnancy timepoints (Mid-pregnancy; Control: 1416 ± 495 vs MNR: 2263 ± 542 ng/mL, p=0.028. Near term; Control: 1784 ± 273 vs MNR: 2587 ± 273 ng/mL, p=0.006).

To determine whether the elevations in circulating cortisol were accompanied by gross morphological changes in the maternal adrenal gland, adrenal morphology was assessed at mid and late gestation (Supplemental Figure 2). There were no significant diet-related differences in total adrenal medulla area, cortex area or medulla-to-cortex ratio at either timepoint (all p > 0.05). There was also no significant diet effects observed in collagen deposition or cell hypertrophy in the zona fasciculata at either timepoint (all p > 0.05).

### Maternal Organ Weights

There was no significant effect of MNR diet on absolute or relative (% body weight) maternal organ weights (heart, lung, liver, kidneys, spleen or ovaries) at mid-pregnancy or near-term (Supplemental Table 3). However, MNR dams showed a trend toward greater absolute liver weight at near-term compared to Controls (Control: 25.42 ± 1.67 vs MNR: 30.27 ± 1.78 g, p = 0.084), which was statistically significant when expressed as a percentage of body weight (Control: 3.21 ± 0.14 vs MNR: 3.81 ± 0.22, p = 0.049). Histological examination of available near-term liver samples revealed apparent tissue lipid accumulation in 4 of 7 MNR dams compared to 1 of 5 Control dams (Supplemental Figure 4).

### Circulating Metabolites

Maternal metabolic profile modestly differed between diet groups, with the most pronounced effects observed at mid-pregnancy (Figure 3). At mid-pregnancy, MNR dams exhibited lower sodium concentrations (Control: 155 ± 8.42 vs. MNR: 130 ± 5.15 mmol/L, p=0.036), lower glucose concentrations (Control: 151 ± 19.5 vs. MNR: 89 ± 18.7 mg/dL, p=0.041) and elevated blood urea nitrogen (Control: 19 ± 0.99 vs. MNR: 24 ± 1.67 mg/dL, p=0.041) compared to Controls, consistent with inadequate energy availability and mobilization of endogenous protein reserves. Additionally, MNR dams showed significantly reduced circulating lipids, including total cholesterol (Control: 32 ± 4.37 vs. MNR: 20.4 ± 0.24 mg/dL, p=0.034), triglycerides (Control: 267 ± 44.9 vs. MNR: 134 ± 15.8 mg/dL, p=0.035), and VLDL (Control: 45.2 ± 4.78 vs. MNR: 26.8 ± 3.15 mg/dL, p=0.021). The acute metabolic perturbations were largely absent near term, though MNR dams showed reduced bicarbonate (Control: 31 ± 0.62 vs. MNR: 29 ± 0.40 mmol/L, p=0.019), suggesting metabolic acidosis. Lipid concentrations remained numerically lower in MNR dams near term but no longer reached statistical significance (all p>0.05).

**Figure 3.**
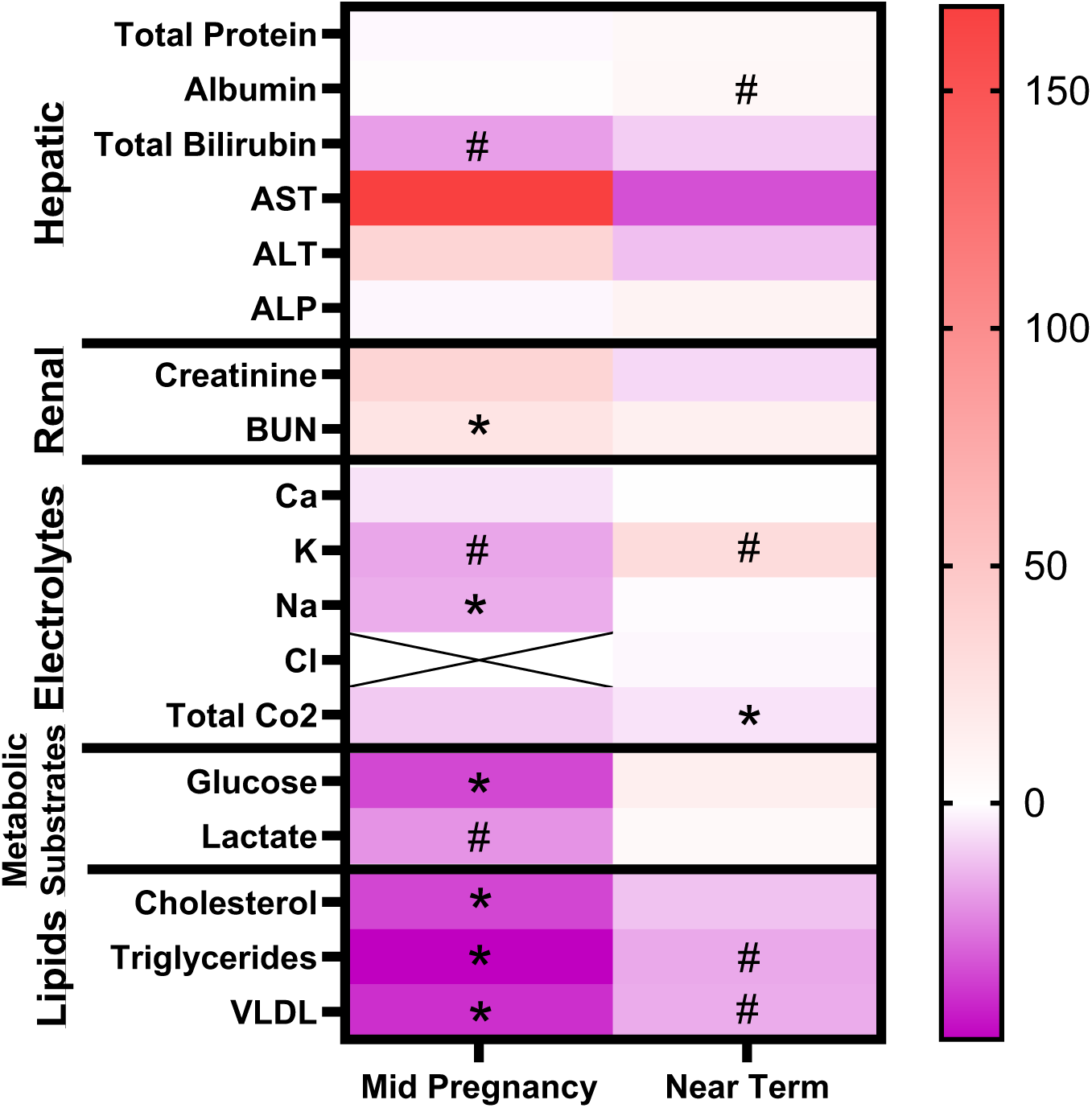
Serum biochemical profiles in maternal nutrient restricted (MNR) dams relative to Control at mid-pregnancy and near term. At mid-pregnancy, MNR dams exhibited lower sodium, glucose and lipids, and elevated blood urea nitrogen compared to Controls. The acute metabolic perturbations were largely absent near term, though MNR dams showed reduced bicarbonate. Lipid concentrations remained numerically lower in MNR dams near term but no longer reached statistical significance. Heat map indicated percent difference in MNR dams compared to Control; red a greater percent difference, purple as reduced percent difference and no difference as white. Mid-pregnancy and near-term analyses were fit separately. #p<0.1; *p<0.05. Mid-pregnancy n=11: 5 Control, 6 MNR. Near-term n=14: 7 Control, 7 MNR

Maternal serum hormone concentrations near term revealed selective rather than global hormonal disruption in MNR dams. Reproductive hormones were unaffected, with no significant differences in progesterone (Control: 29.7 ± 3.70 vs MNR: 33.2 ± 3.71 ng/mL, p>0.05), estriol (Control: 11.1 ± 1.83 vs MNR: 11.3 ± 4.07 pg/mL, p>0.05), DHEA-S (Control: 12.4 ± 1.76 vs MNR: 11.9 ± 0.79 ng/mL, p>0.05) or oxytocin (Control: 15.7 ± 6.39 vs. MNR: 19.2 ± 4.47 pg/mL, p>0.05). MNR dams showed a trend toward elevated serum insulin concentrations (Control: 163 ± 28.7 vs MNR: 234 ± 32.6 pg/mL, p=0.13), although fasting blood glucose concentrations were similar to Control (Control: 234 ± 23.4 vs MNR: 269 ± 32.5 mg/dL, p>0.05).

### Fetal and Maternal-Fetal Interface Outcomes

MNR did not alter the fetal sex ratio. Control dams produced 20 female and 22 male fetuses (47.6% female), while MNR dams produced 17 female and 16 male fetuses (51.5% female; p>0.05), indicating that maternal nutrient restriction did not preferentially affect survival of one sex over the other.

At mid-pregnancy, MNR fetuses showed reduced body weight compared to Controls (p=0.053; Table 1), with the effect most pronounced in female fetuses (−27.6% vs -12.8% in males). MNR fetuses also exhibited a significantly elevated brain:liver weight ratio at mid-pregnancy when compared to Controls (p=0.026), indicative of asymmetric growth restriction with preferential brain growth. Placental weight, fetal:placental weight ratio and fetal circulating glucose did not differ between diet groups (both p>0.05).

**Table 1.** Fetal and Placental Characteristics in Control and maternal nutrient restricted (MNR) dams at mid-pregnancy and near-term.

| *Mid-Pregnancy Fetal and Placental Outcomes* |  |  |  |  |  |  |  |
| --- | --- | --- | --- | --- | --- | --- | --- |
|  | Female Control | Male Control | Female MNR | Male MNR | Female % Diff | Male % Diff | P-value (diet) |
| Fetal Weight | 6.11 ± 0.59 | 5.40 ± 0.64 | 4.42 ± 0.27 | 4.71 ± 0.23 | -27.6 | -12.8 | 0.053 |
| Placental Weight | 3.38 ± 0.23 | 3.34 ± 0.40 | 3.03 ± 0.11 | 2.98 ± 0.12 | NA | NA | 0.220 |
| Fetal:Placental Ratio | 1.81 ± 0.07 | 1.62 ± 0.06 | 1.46 ± 0.10 | 1.58 ± 0.09 | NA | NA | 0.448 |
| Brain:Liver Ratio | 0.75 ± 0.15 | 1.64 ± 0.17 | 1.68 ± 0.24 | 1.90 ± 0.25 | 124.2 | 15.5 | 0.026 |
| Glucose | 61.42 ± 14.87 | 47.25 ± 5.03 | 60.97 ± 5.86 | 53.49 ± 2.93 | NA | NA | 0.929 |
| *Near-Term Fetal and Placental Outcomes* |  |  |  |  |  |  |  |
|  | Female Control | Male Control | Female MNR | Male MNR | Female % Diff | Male % Diff | P-value (diet) |
| Fetal Weight | 79.08 ± 3.37 | 79.31 ± 1.48 | 69.34 ± 3.79 | 71.43 ± 2.48 | -12.3 | -9.9 | 0.019 |
| Placental Weight | 5.33 ± 0.23 | 5.57 ± 0.23 | 5.70 ± 0.33 | 5.83 ± 0.24 | NA | NA | 0.445 |
| Fetal:Placental Ratio | 14.94 ± 0.39 | 14.44 ± 0.61 | 12.46 ± 0.62 | 12.58 ± 0.63 | -16.6 | -12.9 | 0.016 |
| Brain:Liver Ratio | 0.60 ± 0.04 | 0.64 ± 0.03 | 0.67 ± 0.07 | 0.64 ± 0.02 | NA | NA | 0.202 |
| Glucose | 107.82 ± 8.91 | 108.53 ± 9.84 | 101.46 ± 4.92 | 87.13 ± 8.36 | NA | NA | 0.357 |
| Cortisol | 82.22 ± 11.81 | 59.62 ± 8.08 | 147.05 ± 26.34 | 121.85 ± 6.26 | 78.8 | 104.2 | 0.014 |
| Creatine | 11.50 ± 3.18 | 21.38 ± 3.63 | 29.08 ± 5.02 | 30.76 ± 3.42 | 152.8 | 43.8 | 0.018 |

By near-term, MNR fetuses continued to be smaller compared to Control (p=0.019; Table 1) and was associated with reduced fetal:placental weight ratio (p=0.016), indicating disproportionately smaller fetuses relative to placental mass. The brain:liver weight ratio no longer differed significantly between diet groups at term (p=0.202), suggesting resolution or attenuation of the earlier asymmetric growth pattern. However, MNR fetuses showed significantly elevated circulating cortisol (diet effect: p=0.014) and creatine (diet effect: p=0.018) compared to Controls, mirroring the maternal cortisol elevation. Fetal blood glucose concentrations did not differ significantly between diet groups (p=0.357).

## DISCUSSION

Using the well-characterized guinea pig maternal nutrient restriction (MNR) model of FGR, we conducted a comprehensive retrospective analysis to investigate the hypothesis that reduced fetal growth characteristic of maternal undernutrition reflects not only limited nutrient substrate availability, but also the maternal physiological stress response. Our aim was to characterize maternal physiological adaptations across two critical gestational windows and identify stress-related mechanisms contributing to fetal growth restriction beyond direct nutrient limitation. We demonstrated that MNR activated the maternal HPA axis, resulting in sustained elevation of salivary and serum cortisol throughout pregnancy despite no baseline differences between diet groups. The stress response was accompanied by acute metabolic perturbations at mid-pregnancy, including hypoglycemia, hyponatremia, elevated blood urea nitrogen, and depletion of circulating lipids. These changes suggest maternal metabolic stress extending beyond simple substrate limitation. While many acute metabolic changes were resolved by near-term, cortisol elevation remained elevated. MNR fetuses also exhibited elevated circulating cortisol alongside sustained growth restriction. Overall, our data provides evidence that maternal physiological stress responses actively contribute to fetal growth outcomes in undernutrition, shifting the understanding of nutrition-related FGR from a purely substrate-limited model to one in which maternal adaptive capacity and stress physiology become critical determinants of pregnancy outcome.

Despite extensive research on fetal outcomes in maternal undernutrition in the guinea pig^42–49^, few studies have systematically examined the maternal physiological responses to undernutrition during pregnancy. Undernutrition during the first half of pregnancy has been shown to lead to maternal metabolic dysfunction including altered glucose tolerance and insulin sensitivity in rats^50^, supporting the concept that nutritional restriction creates a pathological maternal metabolic state. Our findings extend these observations by characterizing acute metabolic perturbations at mid-pregnancy, including hypoglycemia, hyponatremia, elevated blood urea nitrogen and reduced circulating lipids. Hypoglycemia at this timepoint persisted despite evidence of increased protein catabolism, indicating that compensatory mechanisms were inadequate to maintain glucose homeostasis under the combined metabolic demands of pregnancy and nutrient restriction. This hypothesis warrants further investigation but is beyond the scope of the current study. Additionally, many acute metabolic perturbations observed at mid-pregnancy had largely resolved by near-term. This could be reflective of the methodological transition of the dams from 70% to 90% caloric restriction implemented at GD30 to reduce late pregnancy losses^39^, but likely reflected both increased nutrient availability and potential maternal adaptive mechanisms. Cortisol elevation, however, persisted throughout pregnancy despite the nutritional restoration, indicating sustained HPA axis activation even with improved caloric intake. The selective nature of hormonal disruption, as cortisol remained elevated while reproductive hormones (progesterone, estriol, oxytocin) were unaffected, indicated a targeted stress response.

Sustained elevation of maternal cortisol was the most consistent and pronounced maternal adaptation to chronic nutrient restriction in our study. Notably, cortisol concentrations did not differ between Control and MNR dams at conception, indicating that HPA axis activation was specifically triggered by the physiological demands of pregnancy rather than by dietary restriction alone. This observation is particularly significant given that normal pregnancy is already associated with progressive increases in cortisol production to support maternal metabolic adaptation^51^. The magnitude of cortisol elevation in MNR dams (approximately 2-3-fold increases in both salivary and serum cortisol at mid-pregnancy and near-term) exceeds what is typically observed in the Control pregnancy, suggesting an enhanced stress response to the metabolic challenges of undernutrition. This ongoing HPA axis activation is consistent with elevated maternal glucocorticoids reported in undernourished rats^45, 52^, suggesting a conserved maternal stress response during gestational undernutrition in these species. Validation of cortisol elevation in both saliva and serum strengthens confidence in this finding, as salivary cortisol reflects free, biologically active hormone concentrations^53^ while serum measures represent total cortisol including that bound to binding proteins^54^. Despite sustained cortisol elevation throughout pregnancy, maternal adrenal glands showed no gross morphological changes in total area, cortex-to-medulla ratio, or cellular architecture. This finding contrasts with non-pregnant models where chronic stress typically induces adrenal enlargement and structural remodeling^55, 56^. In normal pregnancy, the maternal adrenal gland undergoes minimal changes, thus it is possible that pregnancy may confer protection against stress-induced morphological changes^57^. The combination of elevated cortisol output without structural adrenal changes indicates enhanced functional capacity rather than tissue remodeling and has important implications for maternal metabolic homeostasis, placental function, and fetal development given cortisol’s wide-ranging effects on glucose metabolism, immune function, and vascular physiology^58^.

Maternal body composition and organ weights revealed key patterns in the response to chronic nutrient restriction that differ from other undernutrition models. Unlike rat models of maternal undernutrition which typically show reduced maternal weight gain throughout pregnancy^59, 60^, MNR guinea pig dams maintained similar weight gain trajectories to controls during early pregnancy. Divergence in maternal weight emerged only after mid-pregnancy, consistent with previous reports utilizing this model^61^. In our study, this late-gestation divergence was primarily driven by reduced conceptus (fetal and placental) weight rather than maternal tissue loss. This pattern suggests successful maternal adaptation to preserve maternal body condition at the expense of fetal growth, consistent with maternal protective mechanisms that prioritize maternal survival and pregnancy completion^62^. Importantly, neither our study nor previous literature observed a reduction in total litter size due to dietary restriction^39, 49, 61, 63^. While prior reports suggested maternal weight at conception had no effect on litter size^61^, our expanded cohort analysis revealed a positive correlation between conception weight and litter size across both diet groups, indicating this model isolates the effects of undernutrition on fetal growth rather than fundamentally altering reproductive capacity. At the organ level, chronic nutrient restriction produced selective rather than generalized effects. The significant increase in relative liver weight in MNR dams at near-term, accompanied by apparent hepatic lipid accumulation, suggests specific alterations in hepatic metabolism. The temporal association between liver enlargement and sustained cortisol elevation is consistent with known effects of glucocorticoids on hepatic metabolism^64, 65^. Importantly, other maternal organs (heart, lung, kidney, spleen, ovary) showed no weight changes, indicating that the hepatic response represented organ-specific adaptation rather than generalized effects of undernutrition.

The parallel elevation of cortisol in MNR mothers and their fetuses represents a critical finding that must be interpreted within the context of our extensive characterization of placental adaptations in this model^66–69^. Our previous work has demonstrated that MNR creates profound placental insufficiency characterized by reduced trophoblast invasion, disorganized labyrinth microvasculature, altered transcriptional programs affecting transmembrane transport and dysregulated growth factor signaling. Importantly, targeted improvement of placental function through IGF1 gene therapy not only rescued fetal growth and corrected these structural and functional placental deficits, but also reduced both maternal and fetal cortisol concentrations at near-term^40^. These results suggest a complex bidirectional relationship where maternal stress physiology may initially compromise placental development during early pregnancy. The resulting placental insufficiency in turn amplifies maternal stress responses, creating a self-reinforcing pathophysiological cycle. The ability of placental-targeted therapy to simultaneously restore placental function, rescue fetal growth, and reduce stress hormone concentrations in both mother and fetus provides compelling evidence that placental health is central to maintaining maternal-fetal homeostasis under nutritional stress. While our current study cannot determine whether fetal cortisol elevation results from direct maternal transfer or fetal HPA axis activation, and is currently under investigation, the therapeutic data suggest that restoring placental function can break the cycle of stress and growth restriction. Overall supporting our hypothesis that maternal stress responses and placental insufficiency are mechanistically linked rather than independent consequences of undernutrition.

Our findings provide mechanistic insights that may explain why nutritional supplementation strategies in pregnancy often show disappointing efficacy in preventing pregnancy complications^70^. The results demonstrate that maternal stress physiology actively mediates fetal growth outcomes beyond simple substrate availability. Therapeutic approaches focused solely on increasing nutrient intake may therefore be insufficient to address the complex pathophysiology underlying nutrition-related growth restriction. However, several study limitations should be acknowledged, including the retrospective design that limits our ability to establish direct causality, modest sample sizes for some analyses and examination of only specific gestational windows that may have missed other critical adaptation periods. The work nevertheless makes valuable contributions to advancing our understanding of FGR pathophysiology. Combined with our extensive placental research demonstrating the interconnection between maternal stress and placental function, these results establish a foundation for future interventional studies targeting the maternal-placental-fetal stress axis. The mechanistic understanding opens new avenues for preventing nutrition-related FGR that extend beyond traditional nutritional approaches and may ultimately lead to more effective interventions for this significant clinical problem.

In conclusion, this study reframes the pathophysiology of nutrition-related FGR, demonstrating that maternal physiological stress in maternal undernutrition is a key mediator of fetal outcomes. Our findings challenge the substrate-limitation paradigm often considered in undernutrition and support a model where maternal HPA axis activation, metabolic reprogramming and placental insufficiency are mechanistically linked. The observation that maternal stress physiology persists even with improved nutritional intake provides a rationale for further studies into why simply providing nutritional supplements may be insufficient to rescue fetal growth. Overall highlighting that effective interventions for FGR in undernourished populations must address both nutritional deficits and the underlying maternal stress response.

## Acknowledgments

We would like to thank veterinary and technical staff at Cincinnati Children’s Hospital and Medical Center and the University of Florida Animal Care Services, and Dr Kendall Stephens, Kristin Lampe, Alyssa Williams, Dr Khanh Huynh, Aditya Mahadevan, Dr Jason Puglise, and Dr Erica Smith for their assistance with animal protocols and necropsies.

## Author Contributions

CR performed experiments, analyzed data and edited manuscript. PR performed experiments, analyzed data and edited manuscript. BND performed animal procedures, and edited manuscript. HNJ obtained funding, supervised and edited the manuscript. RLW conceived the study, obtained funding, supervised, performed experiments, analyzed data, and wrote manuscript. All authors approve the final version of the manuscript.

## Ethics approval

Animal care and usage was approved by the Institutional Animal Care and Use Committees at Cincinnati Children’s Hospital and Medical Center (Protocol number 2017-0065) and University of Florida (Protocol number 202011236).

