## Supplemental Material for "Sustained Maternal Cortisol Elevation Links Metabolic Stress to Fetal Growth Restriction in a Guinea Pig Model of Undernutrition"

**Supplemental Table 1.** Information on sample dilution, and the sensitivity and intra-assay precision reported by the manufacturers for serum assays

| ***Hormone/ Metabolite*** | **Manufacturers Number** | **Serum Dilution** | **Assay Sensitivity** | **Manufacturers Intra-Assay Precision** | **Manufacturers Inter-Assay Precision** |
|---|---|---|---|---|---|
| *Cortisol* | Arbor Assay - K003-H | 1:2000; 1:50 for saliva | 36 pg/mL | 10.1% CV | 12.0% CV |
| *Progesterone* | Arbor Assay - K025-H | 1:50 | 25.3 pg/mL | 3.1-5.1% CV | 4.1-7.0% CV |
| *Estriol* | Arbor Assay - K064-H | Extracted following Steroid Liquid Extraction Protocol using Ethyl Acetate (Arbor Assays) | 5.33 pg/mL | 8.3-13.1% CV | 10.1-17.5% CV |
| *Dehydroepiandrosterone sulfate (DHEA-S)* | Arbor Assay - K054-H | 1:20 | 90.9 pg/mL | 4.6-10.3% CV | 5.2-11.5% CV |
| *Oxytocin* | Arbor Assay - K048-H | Undiluted | 17.0 pg/mL | 4.3-5.2% CV | 7.7-10.0% CV |
| *Insulin* | Novus Biologicals - NBP3-42712 | Undiluted | 51.5 pg/mL | <10% CV | <12% CV |

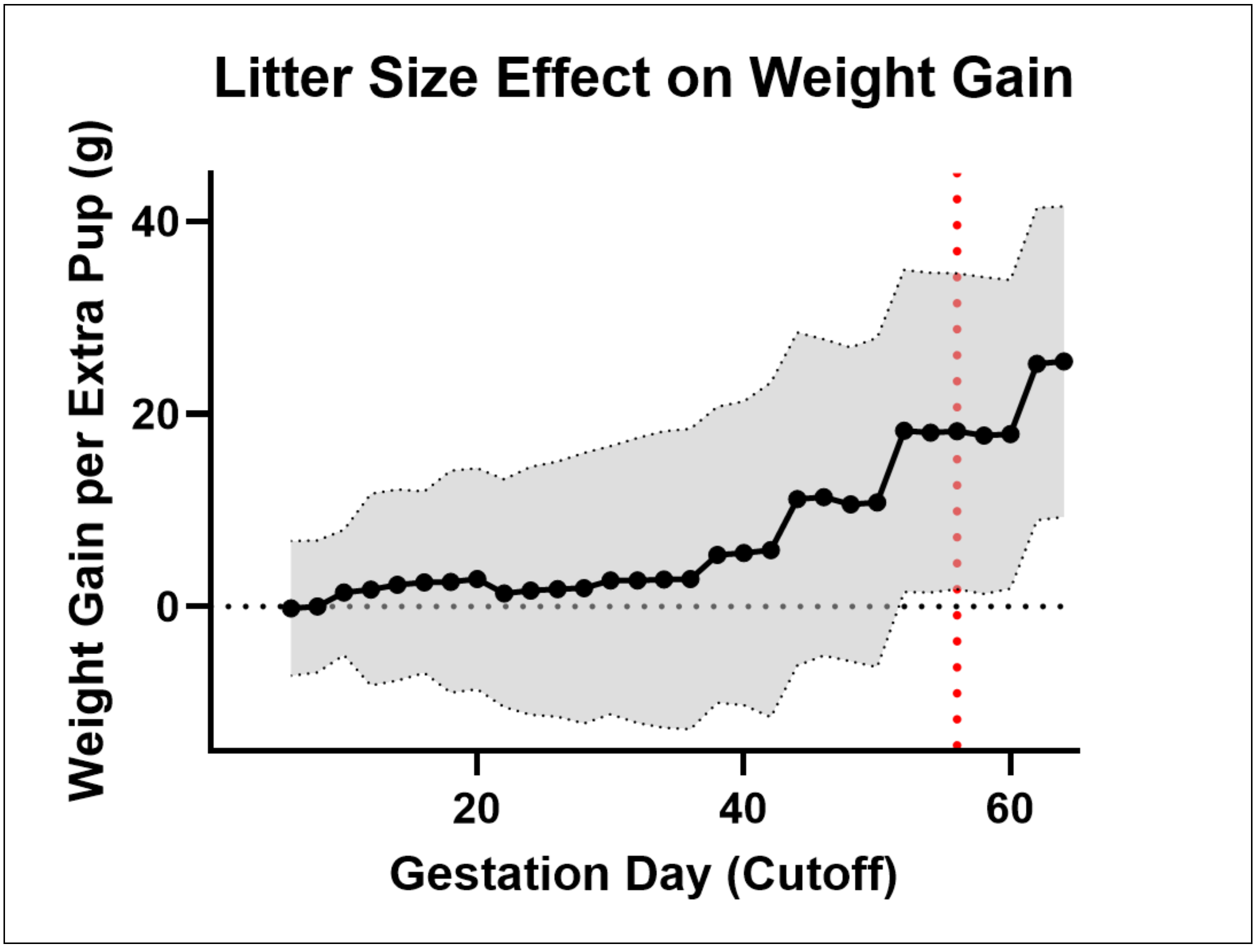


**Supplemental Figure 1. Maternal weight gain in relation to pup number in pregnant guinea pigs.** Sequential analysis revealed that the litter size effect on maternal weight gain emerged progressively during the second half of pregnancy and reached statistical significance by day 56. Over the complete course of pregnancy, each additional pup contributed 25.45 g to total maternal weight gain (95% CI: [9.33, 41.57], $p = 0.008$). Data are the estimated effect of litter size on cumulative weight gain from baseline at each gestational day cutoff (days 0–65, 2-day intervals). Shaded regions represent 95% confidence intervals. The dashed red vertical line indicates the first gestational day at which litter size reached statistical significance ($p < 0.05$). Animals with negative weight gain values were excluded prior to analysis. Data from term cohort only (n=14: 7 control, 7 restricted)

**Supplemental Table 2.** Progressive diet effect on maternal weight gain across pregnancy

| Gestational Week | Control Weight Gain† (g) | Maternal Nutrient Restricted Weight Gain† (g) | Difference‡ (g) | 95% CI | p-value |
|---|---|---|---|---|---|
| 1 | 4.5 ± 9.3 | -7.0 ± 9.1 | 11.5 | [-14.5, 37.5] | 0.389 |
| 2 | 50.5 ± 7.9 | 35.4 ± 7.7 | 15.0 | [-6.9, 36.9] | 0.190 |
| 3 | 96.4 ± 6.8 | 77.9 ± 6.6 | 18.5 | [-1.4, 38.4] | 0.071 |
| 4 | 142.4 ± 6.2 | 120.4 ± 6.0 | 22.1 | [4.8, 39.4] | 0.027 |
| 5 | 188.4 ± 6.3 | 162.8 ± 6.0 | 25.6 | [8.2, 43.0] | 0.013 |
| 6 | 234.4 ± 7.1 | 205.3 ± 6.7 | 29.1 | [9.5, 48.7] | 0.009 |
| 7 | 280.4 ± 8.4 | 247.7 ± 7.9 | 32.6 | [9.8, 55.4] | 0.008 |
| 8 | 326.3 ± 10.0 | 290.2 ± 9.4 | 36.1 | [9.0, 63.2] | 0.011 |
| 9 | 372.3 ± 11.8 | 332.7 ± 11.0 | 39.7 | [7.9, 71.5] | 0.016 |

†Predicted values from mixed-effects model controlling for litter size
‡Control minus Restricted; positive values indicate control animals gained more weight
Data from term cohort only (n=14: 7 Control, 7 MNR)

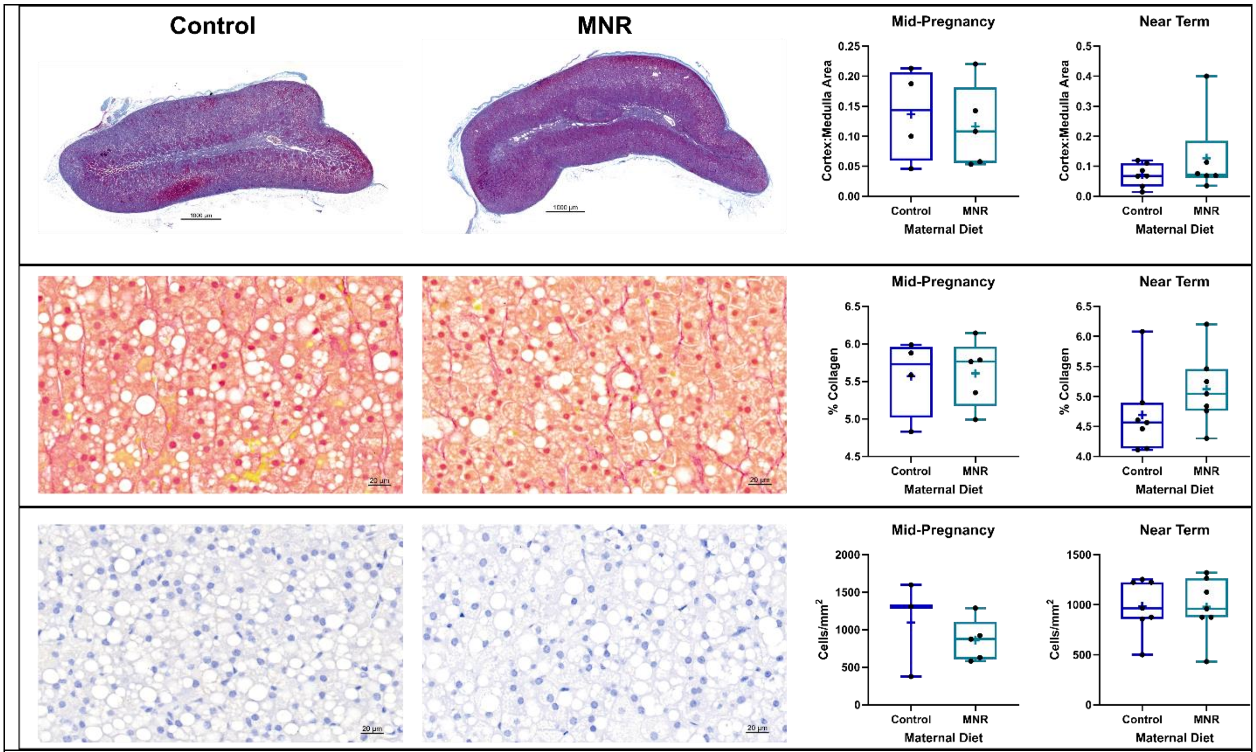


**Supplemental Figure 2. Maternal adrenal gland morphology at mid-pregnancy and near term in Control and maternal nutrient restricted (MNR) dams.** Top panel: There were no significant diet-related differences in total adrenal medulla area, cortex area or medulla-to-cortex ratio at either timepoint. Representative Masson's Trichrome-stained whole adrenal sections from Control and MNR dams. Middle panel: There was also no significant diet effects observed in collagen deposition at either timepoint. Representative Sirius red-stained sections in Control (left) and MNR (right) dams. Bottom panel: There was also no significant diet effects observed in cell hypertrophy in the zona fasciculata at either timepoint. Representative hematoxylin stained sections in Control (left) and MNR (right) dams. Data is median and interquartile range; crosses (+) indicate group means; individual points are overlaid. n=9: 4 Control, 5 MNR

**Supplemental Table 3.** Impact of maternal nutrient restriction (MNR) on maternal organ weights at two pregnancy timepoints

| | | Absolute Weight | | | % Carcass Weight | | |
|---|---|---|---|---|---|---|---|
| Timepoint | Organ | Control (g) | MNR (g) | P-value | Control (% carcass) | MNR (% carcass) | P-value |
| Mid-pregnancy | Heart | 3.63 ± 0.26 | 3.20 ± 0.19 | 0.253 | 0.46 ± 0.04 | 0.48 ± 0.02 | 0.686 |
| | Lung | 5.60 ± 0.31 | 6.27 ± 0.98 | 0.559 | 0.72 ± 0.05 | 0.97 ± 0.17 | 0.187 |
| | Liver | 27.47 ± 2.13 | 22.79 ± 1.33 | 0.348 | 3.44 ± 0.12 | 3.44 ± 0.12 | 0.919 |
| | Kidney | 6.09 ± 0.30 | 5.08 ± 0.36 | 0.267 | 0.77 ± 0.03 | 0.76 ± 0.01 | 0.911 |
| | Spleen | 0.85 ± 0.08 | 0.61 ± 0.06 | 0.238 | 0.11 ± 0.01 | 0.09 ± 0.01 | 0.332 |
| | Ovary | 0.14 ± 0.02 | 0.18 ± 0.06 | 0.479 | 0.02 ± 0.00 | 0.03 ± 0.01 | 0.231 |
| Near-term | Heart | 3.16 ± 0.09 | 2.96 ± 0.23 | 0.461 | 0.40 ± 0.02 | 0.37 ± 0.03 | 0.374 |
| | Lung | 5.37 ± 0.35 | 5.36 ± 0.48 | 0.947 | 0.68 ± 0.04 | 0.67 ± 0.05 | 0.846 |
| | **Liver** | **25.42 ± 1.67** | **30.27 ± 1.78** | **0.084** | **3.21 ± 0.14** | **3.81 ± 0.22** | **0.049** |
| | Kidney | 5.75 ± 0.29 | 6.19 ± 0.21 | 0.264 | 0.73 ± 0.03 | 0.78 ± 0.04 | 0.292 |
| | Spleen | 0.89 ± 0.05 | 0.86 ± 0.07 | 0.735 | 0.11 ± 0.01 | 0.11 ± 0.01 | 0.576 |
| | Ovary | 0.15 ± 0.02 | 0.11 ± 0.02 | 0.155 | 0.02 ± 0.00 | 0.01 ± 0.00 | 0.130 |

Data are mean ± SEM. Carcass weight was calculated as total maternal weight minus conceptus (fetus and placenta) weight. Mid-pregnancy n=11: 5 Control, 6 MNR. Near-term n=14: 7 Control, 7 MNR

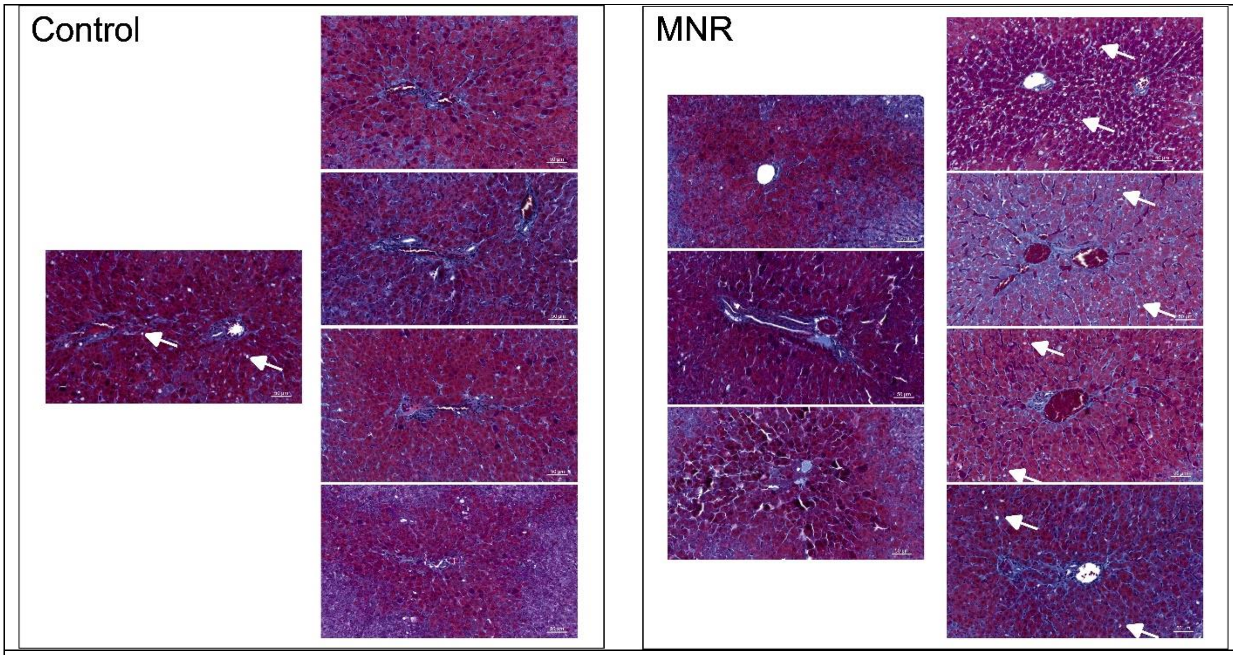


**Supplemental Figure 4. Histological examination of available near-term liver samples in Control and maternal nutrient restricted (MNR) dams**. Apparent tissue lipid accumulation was observed in 4 of 7 MNR dams compared to 1 of 5 Control dams. White arrows indicate lipid droplet.
